# Hippocampal ripple-mediated convergence of spatial codes drives the emergence of cortical contextual representations

**DOI:** 10.1101/2021.03.29.437441

**Authors:** Ayaka Bota, Yuki K. Murai, Xinzhi Jiang, Suzune Tsukamoto, Alessandro Luchetti, Akihiro Goto, Toru Takumi, Steven J. Middleton, Yasunori Hayashi

**Affiliations:** Graduate School of Science and Engineering, Saitama University, Saitama 338-8570, Japan; Department of Pharmacology, Kyoto University Graduate School of Medicine, Kyoto 606-8501, Japan; Brain Science Institute, RIKEN, Wako, Saitama 351-0198, Japan; Department of Physiology and Cell Biology, Kobe University School of Medicine, Kobe 650-0017, Japan; Brain and Body System Science Institute, Saitama University, Saitama 338-8570, Japan

**Keywords:** Spatial context cells, sharp-wave ripples, replay, anterior cingulate cortex, hippocampus, place cells, memory consolidation

## Abstract

Hippocampal sharp-wave ripples (SWRs) and the associated replay of neuronal sequences are well-established as the primary physiological substrates for memory consolidation. While the local transformations they induce within the hippocampus are well-characterized, their impact on the downstream cortical representations remains unknown. Here, we identify a distinct functional class of neurons in the anterior cingulate cortex “spatial context cells” that emerge through the multi-day integration of discrete hippocampal inputs. Using closed-loop disruption of SWRs and longitudinal calcium imaging, we demonstrate that formation of this global spatial framework depends on SWRs during offline periods. Whereas standard cortical place maps remain intact following SWR disruption, the emergence of spatial context cells is selectively abolished. These cells preferentially express c-Fos, identifying them as constituents of the cortical memory engram, and exhibit broad receptive fields that develop through an experience-dependent process of field convergence. Collectively, these findings establish SWR-mediated replay as a computational bridge transforming high-resolution, fragmented hippocampal spatial codes into generalized, stable cortical schemas. This work reveals a bifurcated consolidation process, identifying SWR-dependent synaptic remodeling as a fundamental mechanism for the ontogeny of remote memory.

## Main text

The stabilization of episodic memories relies on a systems-level dialogue between the hippocampus and the neocortex. Central to this process are hippocampal sharp-wave ripples (SWRs), high-frequency oscillations that occur during sleep and quiet wakefulness. These transient events are known to temporally compress and replay sequences of neuronal firing that occurred during awake periods ^1,2^, a mechanism theorized to be the essential driver for memory consolidation ^3,4^. While the local consequences of SWRs within the hippocampus have been characterized extensively ^5,6^, demonstrating that these events reshape local ensemble dynamics to refine internal spatial representations^7^, most current models are limited by a hippocampal-centric perspective. Although we recognize SWRs as essential for long-term memory, the precise downstream mechanisms by which they broadcast and transform information to the neocortex, thereby shaping cortical representations, remain poorly understood. This gap in our knowledge, the lack of a cellular-level description of how hippocampal replay sculpts cortical circuitry, is a significant barrier to understanding systems consolidation. While prevailing theories posit that the neocortex synthesizes generalized environmental “schemas” from hippocampal experience ^8,9^, it remains unclear whether cortical representations are merely passive reflections of hippocampal place fields or unique functional entities driven by SWR-dependent plasticity. Understanding how the discrete, high-resolution hippocampal “pixels” of space are transformed into the unified, large-scale contextual representations that characterize remote memories is essential to resolving how the cortex transcends its role as a passive repository to become the active orchestrator of remote, contextual memory.

To address this, we employed a closed-loop system for the real-time disruption of offline SWRs during sleep and quiet wakefulness^3,6,10^, coupled with longitudinal calcium imaging in the anterior cingulate cortex (ACC), a region critical for long-term memory retrieval^11^. This identified a previously undescribed population of excitatory ACC neurons, termed “spatial context cells,” which failed to emerge when SWRs were disrupted. These neurons were characterized by exceptionally broad receptive fields that formed dynamically across the course of learning, in a timeline consistent with memory consolidation and were preferentially recruited into the *c-Fos*-defined memory engram, establishing their role as a substrate for remote memory. Crucially, our experiments reveal a bifurcated consolidation process: while basic cortical spatial tuning is unaffected by SWR disruption, the emergence of spatial context cells is selectively abolished. These data indicate that SWRs function as a computational bridge, driving the synaptic convergence of fragmented hippocampal place codes into a generalized cortical contextual framework. These findings provide a mechanistic foundation for the emergence of environmental contextual representations, identifying SWRs not merely as markers of hippocampal activity, but as the active drivers of downstream cortical synthesis.

## Results

### Disruption of hippocampal SWRs decreases specific neuronal subpopulation in the ACC

Neuronal activity was imaged from ACC layer 2/3 in mice expressing G-CaMP7 under the control of the CaMKIIα promoter during a spatial navigation task **(Fig. 1a, b and Extended Data Fig. 1)**. Mice navigated a sequential behavioral protocol consisting of a square track (S), followed by an open field (O) before being returned to the square track (S’), each for 10 minutes, repeated for nine days **(Fig. 1c)**. ACC neurons exhibited canonical place cell properties, including location specificity, reproducibility across laps, and running direction selectivity, similar to hippocampal CA1 pyramidal cells (**Fig. 1d and Extended Data Fig. 1)** ^12,13^. To characterize differences in spatial representation between ACC and CA1, we applied t-distributed stochastic neighbor embedding (t-SNE), a method for nonlinear dimensionality reduction, to classify the activity of all recorded neurons during the square track sessions (S and S’) across all nine days. When the resulting two-dimensional projections were color-coded by place field position, the maps revealed a similarity in how both brain regions represent spatial information (**Fig. 1e and Extended Data Fig. 1**).

**Figure 1.**
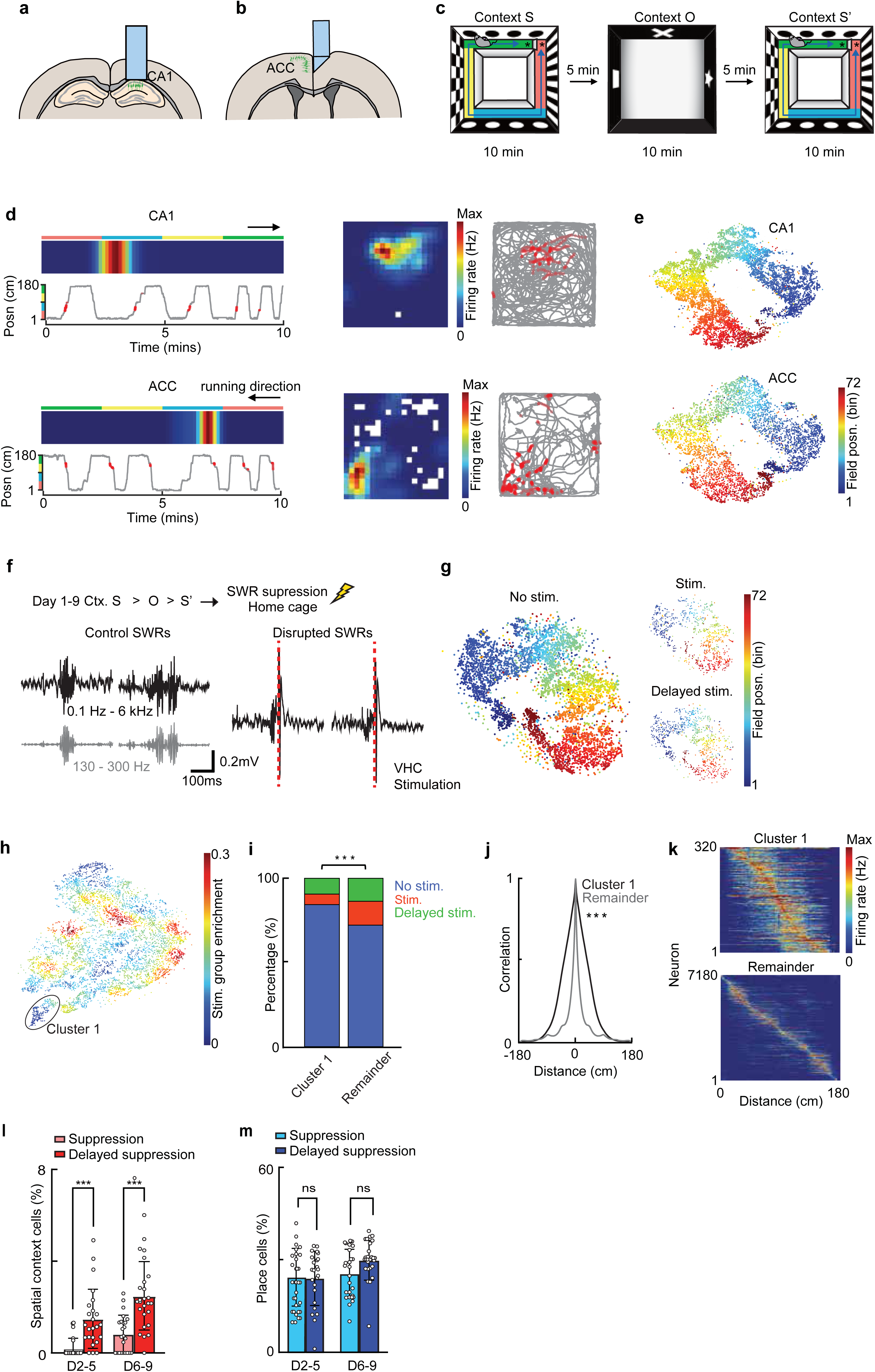
Sharp-wave ripples are required for the formation of spatial context representations in ACC. (**a**) Schematic showing hippocampal imaging strategy, with GRIN lens implanted above the CA1 cell body layer. (**b**) Same as in (a) but for ACC, using a microprism inserted into the fissure to image ACC. (**c**) Overview of the experimental protocol, depicting the square track (contexts S and S’) and open field (context O), reward locations in S are indicated by asterisks. (**d**) Representative examples of neurons imaged from CA1 (top) and ACC (bottom) showing spatial tuning in the two different contexts. Heatmaps depict occupancy normalized firing ratemaps, with corresponding behavioral trajectories shown in gray (binned position) and position of calcium events overlaid in red. (**e**) Combined t-SNE clustering of each neuron’s activity in the S & S’ sessions, from both dCA1 (top) and ACC (bottom), each dot is color-coded to mean place field position on the square track. 1946 unique ACC neurons (from 4 animals, totaling 5453 neurons across all sessions) and 3458 unique dCA1 neurons (from 5 animals, totaling 9150 neurons across all sessions). (**f**) Sharp-wave ripple disruption strategy. (**g**) t-SNE clustering of ACC ratemaps, for control (left), SWR disruption (top right) and delayed stimulation groups (bottom right), each dot is color-coded to mean place field position. (**h**) t-SNE clustering of all ACC neurons auto-correlated ratemaps. Neurons are color coded according to the enrichment of SWR disruption group in the surrounding population. The black circle encapsulates a population separated from the main body of neurons. (**i**) Comparison of cluster composition across experimental groups. The target cluster (Cluster 1) was significantly depleted of neurons from the SWR-disruption group compared to control and delayed-stimulation cohorts (χ2 = 24.28, d.f. = 2, P = 5.3 × 10-6, χ2 contingency test, n = 320 neurons in cluster 1, n = 7180 neurons in the remainder. Data are presented as the observed versus expected distribution of neurons by group. (**j**) Mean auto-correlograms of all neurons contained within the main population and cluster 1 (Half-width, P = 1.5 × 10-197, Mann-Whitney U test). (**k**) The sorted ratemaps of all ACC neurons in the two populations. (**l**) Fraction of spatial context cells in the square track during days 2-5(early) and 6-9(late), for the suppression (pink) and delayed-suppression (red) groups. A two-way repeated-measures ANOVA revealed significant main effects of time (F(1,46) = 18.87, P < 0.0001) and group (F(1,46) = 33.88, P < 0.0001), but no significant interaction (F(1,46) = 1.00, P = 0.3224). Post hoc comparisons using Fisher’s LSD test showed that the fraction of SC cells was significantly lower in the suppression group compared to the delayed-suppression group in both the early (P < 0.0001) and late (P < 0.0001) phases. (**m**) Same as in (l), but for ACC place cells. A two-way repeated-measures ANOVA revealed a significant main effect of time (F(1,46) = 9.52, P = 0.0034), but no significant effect of group (F(1,46) = 0.91, P = 0.3439). A small but significant interaction was observed (F(1,46) = 4.50, P = 0.0394). Post hoc comparisons using Fisher’s LSD test showed no significant difference between suppression and control groups in either the early (P = 0.8718) or late (P = 0.0680) phases.

To address how offline SWRs influence the representation within the extra-hippocampal circuits, we employed a closed-loop system for real-time suppression of hippocampal SWRs, delivering electrical stimulation to the ventral hippocampal commissure upon SWR detection^3,6,10^. After the S-O-S’ session, we returned the animals to their home cage and disrupted SWRs using the system for 1 hour across nine consecutive days (**Fig. 1f**). A control cohort received delayed stimulation (200 ms post-detection), with both groups undergoing the protocol for one hour daily following spatial exploration across nine consecutive days. Initial t-SNE visualization of ACC activity revealed no gross structural reorganization of spatial representations following SWR disruption (**Fig. 1g**). To isolate functional dynamics from spatial topology, we performed a manifold analysis on auto-correlated rate maps, effectively removing the spatial component. By color-coding neurons based on their local density within the SWR-disruption population, we identified a distinct sub-cluster (Cluster 1) that was markedly depleted of neurons from the SWR-disrupted group (**Fig. 1h**). Quantification of cluster composition confirmed that the fraction of SWR-disrupted neurons within Cluster 1 was significantly lower than predicted by chance (**Fig. 1i**) (*P* = 5.3 × 10^−6^, χ^2^ contingency test). Further, analysis of mean auto-correlograms revealed a sharp functional divergence between populations, with the majority of neurons exhibiting standard spatial tuning, Cluster 1 neurons maintained high correlations across significantly larger spatial lags (*P* = 1.5 × 10^−197^, Mann-Whitney *U* test; **Fig. 1j**). This suggested that SWRs are specifically required for the emergence of a sub-population of ACC neurons characterized by exceptionally large spatial receptive fields (**Fig. 1k**). By plotting field-size distributions, we determined that a 62.5 cm threshold optimally captured the majority of these broadly tuned neurons (**Extended Data Fig. 2**). Projecting this population onto the initial t-SNE manifold confirmed that these neurons constitute a distinct functional cluster within the ACC spatial map (**Extended Data Fig. 2**). Further, our analysis revealed that the fraction of these neurons increased significantly with experience (days 1–3: 6.4 ± 0.9% vs 14.3 ± 2.3% on days 7-9, two-way ANOVA, *p* < 0.0001; **Extended Data Fig. 2**) specifically in ACC, but not in dCA1 (days 1-3: 0.3 ± 0.1%; days 7-9: 0.3 ± 0.1%). The emergence of cells with large spatial receptive fields was selectively abolished in the disruption group (**Fig. 1l**), whereas canonical ACC place cells developed normally (**Fig. 1m**).

In order to confirm the selective requirement of hippocampal offline neuronal activity, we additionally used chemogenetics. We expressed inhibitory hM4Di in hippocampus bilaterally and administered a DREADD agonist C21 after the S-O-S’ sessions, each day (**Fig. 2a**). The C21 group exhibited a significantly smaller fraction of cells with large spatial receptive field compared to controls (**Fig. 2b**), consistent with the result of SWR electrical suppression. In contrast, offline C21 administration did not alter the proportion of place cells (**Figs. 2c**), though it did reduce their reliability (**Fig. 2d**).

**Figure 2.**
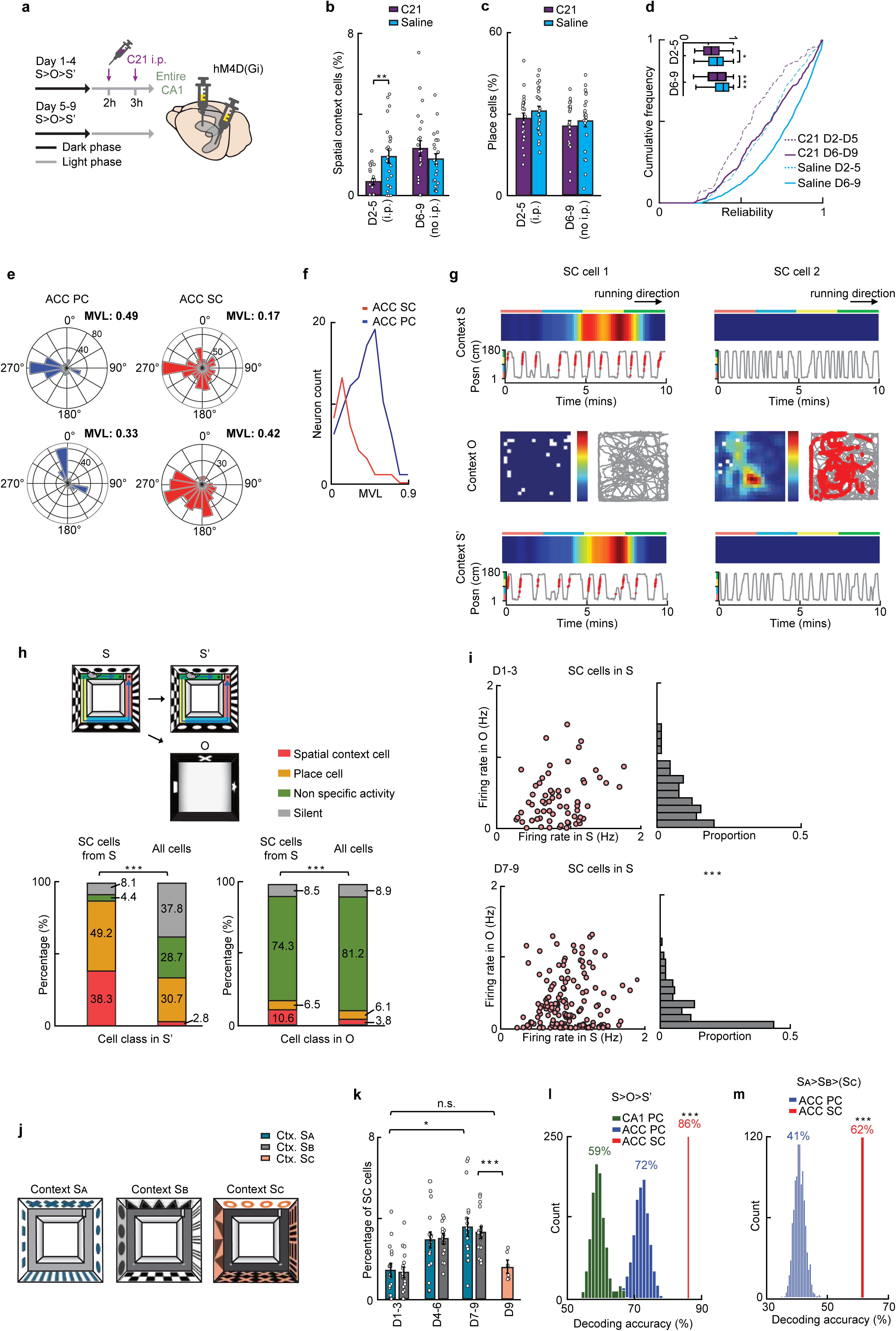
ACC large place field neurons fire in a context selective manner. (**a**) Experimental protocol. (**b**) Proportion of spatial context cells across days 2–5 and 6–9 in the C21 (violet) and saline (blue) groups. Mixed-effects model (REML) revealed a significant main effect of time (F(1, 42) = 6.58, P = 0.014) and a significant interaction between time and group (F(1, 42) = 9.013, P = 0.0045), whereas the main effect of group was not significant (F(1, 46) = 1.47, P = 0.23). Post hoc Šidák’s multiple comparisons test showed a significant difference between groups during days 2–5 (P = 0.0082), but not during days 6–9 (P = 0.42). (**c**) Fraction of place cells across days 2–5 and 6–9 in the C21 (violet) and saline (blue) groups. Two-way repeated-measures ANOVA test revealed a significant main effect of time (F(1, 44) = 6.82, P = 0.012), but no significant main effect of group (F(1, 44) = 0.55, P = 0.46) or interaction (F(1, 44) = 0.0043, P = 0.84). Post hoc Fisher’s LSD test showed no significant differences between groups before (P = 0.64) or after C21 treatment (P = 0.47). (**d**) Cumulative frequency distribution of reliability of ACC place cells during days 2–5 (D1–4, suppression period) and days 6–9 (D5–9, after suppression) in the C21 (violet) and saline (blue) groups. Two-way ANOVA test revealed a significant main effect of group (F(3, 899) = 21.13, P < 0.0001), but no significant time factor effect (F(819, 899) = 1.057, P = 0.2066). Post hoc Tukey’s test showed significant differences between groups during both D2–5 (P = 0.014) and D6–9 (P <0.0001). (**e**) Representative examples of head direction tuning for two ACC place cells (left, blue) and two spatial context cells (middle, red). **(f)** Mean vector lengths (MVL) for all neurons on day 9. (**g**) Representative activity of two ACC spatial context cells (SC cells) from one session across the three contexts. **(h)** Activity classification of SC cells identified in S, in S′ (n = 159 neurons from 4 mice, D7–9) compared with overall distribution in S′ (left), and in O compared with the overall O distribution (right). χ² tests, S vs. S′: p = 8.1 × 10⁻¹⁴⁶; S vs. O: p = 0.00041. **(i)** Firing rates of spatial context cells in S across S and O early (D1–3) and late (D7–9) sessions. Each dot represents a single neuron (early: 59 neurons from 24 sessions from 4 mice; late: 145 neurons from 24 sessions from 4 mice). The histograms on the right show the normalized distributions of O firing rates, which significantly differed between early and late sessions (Kolmogorov–Smirnov test, *p* = 0.0045, KS = 0.26). **(j)** Experimental protocol. Mice were exposed to contexts S_A_ and S_B_ for 9 days, and S_C_ only on day 9. **(k)** The proportion of spatial context cells increased from early (D1–3; n = 18 sessions from 3 mice) to late (D7–9; n = 18 sessions) sessions in both S_A_ (p = 0.0025) and S_B_ (p = 0.0039), (D1–3 vs. D4–6, n.s.). On day 9, S_C_ (D9; n = 6 sessions from 3 mice) showed a significantly lower proportion of SC cells compared with S_A_ (p = 0.0046) and S_B_ (p =0.035). Mixed-effects model with Holm-adjusted p values. **(l)** Decoding accuracy of context identity (S vs. O) using ACC spatial context (SC) cells (red), compared with distributions obtained from equal numbers of CA1 place cells (green) or ACC place cells (blue) randomly selected 1000 times. **(m)** Decoding accuracy of context identity (S_A_, S_B_, S_C_) using ACC spatial context cells (red, 62%) compared to ACC place cells (blue, mean 41%; n = 93 cells from 3 mice). Two-tailed permutation tests with 1000 shuffles.

Postsubiculum neurons in the parahippocampal formation show selective tuning to head direction ^14^. We wondered if this feature is also observed in neurons in ACC. We quantified the degree to which ACC neurons exhibited head-direction tuning as mice explored context O freely on day 9 **(Extended Data Fig. 3)**. The degree to which ACC neurons with broad spatial fields were tuned to head direction changes was significantly weaker compared to typical ACC place cells (p<0.0001, Ranksum, N=4, 4 n=38, 94) (**Fig. 2e and 2f**).

These data suggest that the ACC harbors a functionally distinct neuronal subpopulation which we term spatial context cells, characterized by spatial receptive fields that span significantly broader areas than place cells in the same area as well as head-direction tuning. We propose that while the hippocampus maps discrete local features, these ACC cells emerge through a process of SWR-dependent consolidation during offline periods to integrate fragmented spatial inputs into a unified representation of the global environment. Different sensitivity of spatial context cells and place cells to SWR suppression, by either electrical or chemogenetic approaches, suggests a bifurcated consolidation process, where SWRs are not required for the maintenance of basic spatial tuning in the cortex, but are indispensable for the plasticity-driven synthesis of large-scale contextual representations through convergent hippocampal input.

### Experience-dependent formation of spatial context cells of a discriminative, cortical code for environmental identity

To characterize these cells further, we analyzed their activity across the three contexts (S, O, and S’) during the late training phase (Days 7–9) (**Fig. 2h and Extended Data Fig. 2**). Among cells identified as spatial context cells in context S, a large proportion (38.3 ± 5.3%) maintained their large field in the subsequent S’ session (**Fig. 2h**). An additional 49.2 ± 4.8% of these cells transitioned to conventional place cells in S’, often with receptive fields partially overlapping the original S field (**Extended Data Fig. 2**). In contrast, in the distinct open field context (O), only 10.6 ± 3.0% of the original S spatial context cells maintained their large fields, a significantly smaller fraction than in S’. The remaining neurons either became place cells (6.5 ± 2.1%), exhibited unorganized firing (74.3 ±4.1%), or were silent (8.5 ± 2.1%) in context O. Critically, the overall number of spatial context cells detected in each environment (S and O) was comparable (**Extended Data Fig. 2**), demonstrating that these two distinct contexts recruit largely non-overlapping populations of spatial context cells. Despite the high degree of remapping, approximately 10% of the spatial context neurons were common to both environments, suggesting that while most cells are context-specific, a small subset may encode the sequence or association between the environments due to the multiple daily sequential (S→O→S’) exposures.

To investigate how learning modulates the specificity of spatial context cells, their firing rates (within a single day) were compared between contexts S and O during early (Days 1–3) and late (Days 7–9) sessions (**Fig. 2i**). There was no significant correlation between activity in S and O at either time point (Spearman’s rank correlation: early, *r*=0.25, *p*=0.10; late, *r*=0.04, *p*=0.62). Importantly, activity of spatial context cells in S was reduced in O in later sessions compared with earlier sessions, indicating an increased specificity as animals became more familiar with the context (Kolmogorov–Smirnov test, *p* = 0.0045, KS = 0.26).

To confirm that contextual specificity is a fundamental property of spatial context cells that differentiates them from other spatially selective neurons, we employed three square environments (S_A_, S_B_, S_C_) that differed only in sensory features (see Methods). Mice were trained in S_A_ and S_B_ for 8 days, with only a single exposure to S_C_ on Day 9 (**Fig. 2j**). S_A_ and S_B_ were associated with a similar proportion of spatial context cells (3.5±0.4% and 3.3±0.2%, respectively), but the proportion for the novel S_C_ environment was significantly lower (1.6±0.3%; Holm-adjusted p<0.01 for S_A_-late vs. S_C_ and S_B_-late vs. S_C_; n=3 mice) (**Fig. 2k**). This indicates that spatial context cells emerge in a context-dependent manner requiring experience, rather than forming non-specifically to any environment.

To test our hypothesis that spatial context cells preferentially encode information about the context in which an animal is currently in, rather than its absolute position, we used a support vector machine-linear classifier to test if the identity of the environment could be decoded from the neuronal activity (**Fig. 2l**). The model, trained on 20% of spatial context cell data from S and O, predicted the environment identity with 86% accuracy. This performance was significantly superior to decoding using an equivalent number of randomly selected hippocampal or ACC place cells (Two-tailed permutation test, 1000 shuffles, p<0.0001 vs. both control groups). Interestingly, ACC place cells also decoded context better than hippocampal place cells, underscoring both the ability of spatial context cells to strongly discriminate between environments (**Fig. 2l**), but also indicating that context specificity is an intrinsic feature of the ACC spatial code. Decoding environment identity from the three perceptually similar contexts (S_A_–S_C_) yielded complementary results, with ACC spatial context cells demonstrating significantly better discrimination (62% accuracy) compared to ACC place cells (41%) (**Fig. 2m**).

### Spatial context cells emerge as a functionally distinct class via the integration of fragmented spatial inputs

Spatial context cells emerge dynamically as familiarity with an environment increases (**Fig. 4a, b, 2d, Extended Data Fig. 2**). To better understand this process, we retrospectively and prospectively tracked the activity of neurons from the session in which they first met the spatial context cell criteria. In many cases spatial context cells developed from neurons previously exhibiting place cell-like activity that expanded as a function of experience to form a larger field (**Fig. 3a**). Specifically, on the preceding day, 43% (114 of 268 total cells) were classified as place cells, while the remainder (57%, 154 cells) emerged without prior place cell-like activity (**Fig. 3b**). Intriguingly, unlike dCA1 place cells, which typically had a single field (83%), a high proportion of ACC place cells that later transitioned to spatial context cells had multiple fields (68% with ≥2 fields; 32% with a single field) (**Fig. 3b**). Analyzing all ACC place cells, regardless of their later fate, showed a lower proportion with multiple fields (39% multiple vs. 61% single for D7-D9) (**Extended Data Fig. 3**), suggesting that multiple place fields may be an intrinsic precursor property distinguishing ACC place cells with the potential to converge into a larger spatial context field. Finally, once formed, the spatial representation of a spatial context cell was on average, more stable than both hippocampal and ACC place cells (**Extended Data Fig. 3**). The experience-dependent expansion of this population specifically within the ACC supports a model where the cortex progressively extracts a stable ‘contextual’ framework of a familiar environment from the transient, high-resolution spatial streams broadcast by the hippocampus.

**Figure 3.**
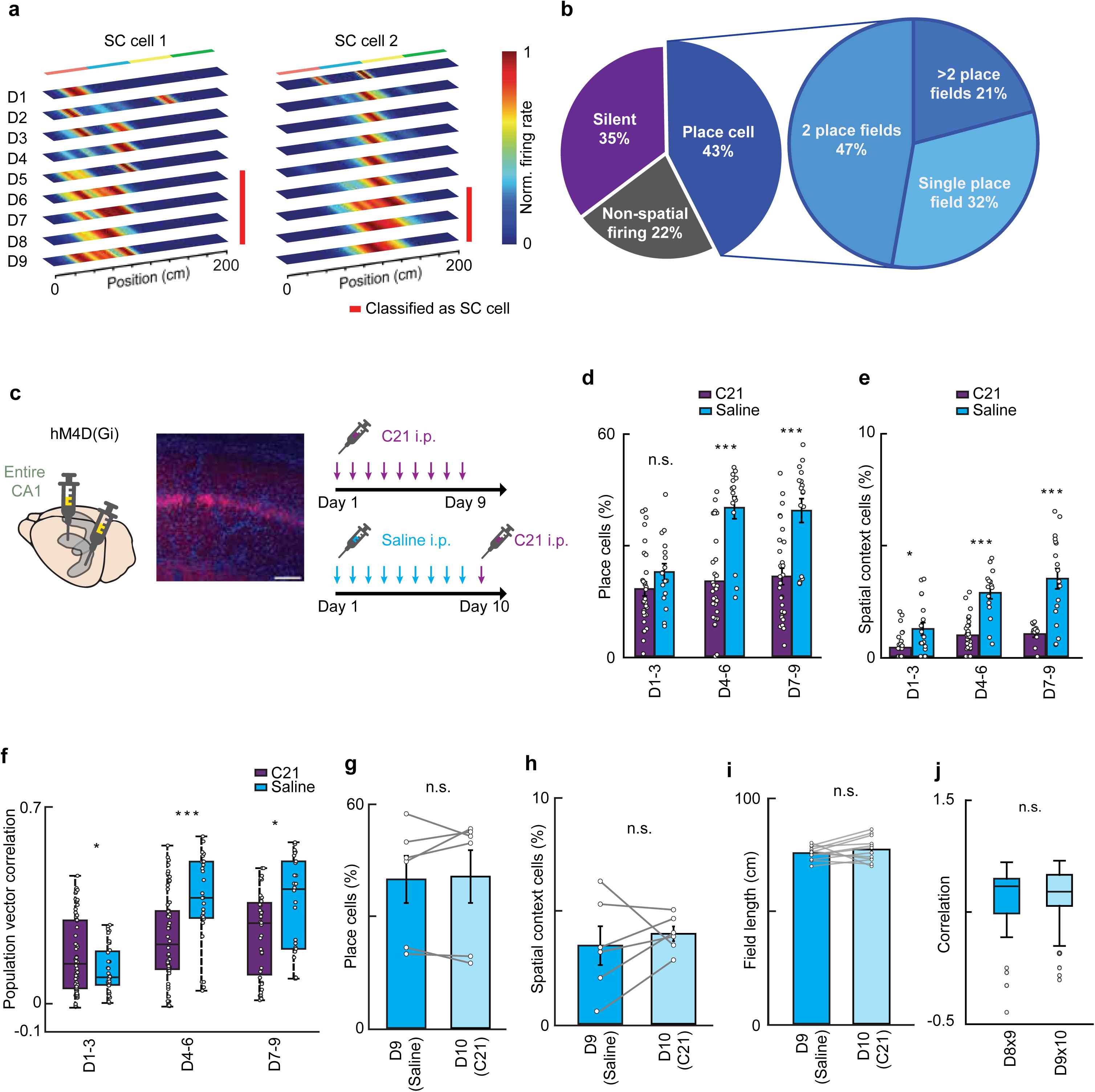
Hippocampal activity is required for the initial formation of spatial context cells. (**a**) Representative examples of neurons that were later classified as SC cells (red bar), tracked across all days (D1–9). (**b**) Activity profile of SC cells on the day prior to first being classified as SC cells (n = 268 neurons from 4 mice, D1–9, left). The number of place fields in the fraction of neurons that were classified as place cells on the prior day (right). (**c**) Experimental protocol for chemogenetic suppression of hippocampal activity. Scale bar =100μm (**d**) Proportion of place cells in the ACC in the saline and C21 groups (session-level data). Mixed-effects model revealed significant main effects of time (F(1.762, 79.30) = 23.16, P < 0.0001) and group (F(1, 46) = 20.36, P < 0.0001), as well as a significant interaction (F(1.762, 79.30) = 12.12, P < 0.0001). Post hoc Šidák’s test indicated no difference between groups in days 1–3 (P = 0.32), but a significant reduction in the C21 group in days 4–6 (P = 0.0001) and 7–9 (P = 0.0006). **(e)** Proportion of spatial context cells ACC in the saline and C21 groups (session-level data). Mixed-effects model (Restricted Maximum Likelihood) revealed significant main effects of time (F(1.423, 64.04) = 28.29, P < 0.0001) and group (F(1, 46) = 62.58, P < 0.0001), as well as a significant interaction (F(1.423, 64.04) = 8.808, P = 0.0015). Post hoc Šidák’s test indicated that the proportion of spatial context cells was lower in the C21 group than in the Saline group during days 1–3 (P = 0.038), 4–6 (P < 0.0001), and 7–9 (P = 0.0001). (**f**) Across-day population vector correlation of ACC neurons. Mixed-effects model revealed significant main effects of time (F(1.991, 183.1) = 56.86, P < 0.0001) and group (F(1, 118) = 13.82, P = 0.0003), as well as a significant interaction (F(1.991, 183.1) = 17.98, P < 0.0001). Post hoc Šidák’s test indicated that correlations were lower in the C21 group than in the Saline group during days 1–3 (P = 0.048), 4–6 (P < 0.0001), and 7–9 (P = 0.0029). Data are shown as box-and-whisker plots (center line, median; box, interquartile range; whiskers, 1.5 × interquartile range). **(g)** Percentage of ACC place cells on days 9 (saline) and 10 (C21) (P = 0.7468, paired t-test). (**h**) Percentage of spatial context cells on days 9 (saline) and 10 (C21) (P = 0.0887, paired t-test). (**i**) Place field length of spatial context cells on days 9 (Saline) and 10 (C21) (6 sessions per day, 3 mice) (P = 0.30, paired t-test). **(j)** Across-day rate-map correlations of spatial context cells, between days 8&9 (n = 391 neurons) and days 9&10 (n = 343 neurons, t(431.5) = 1.129, P = 0.26, Welch’s t-test).

**Figure 4.**
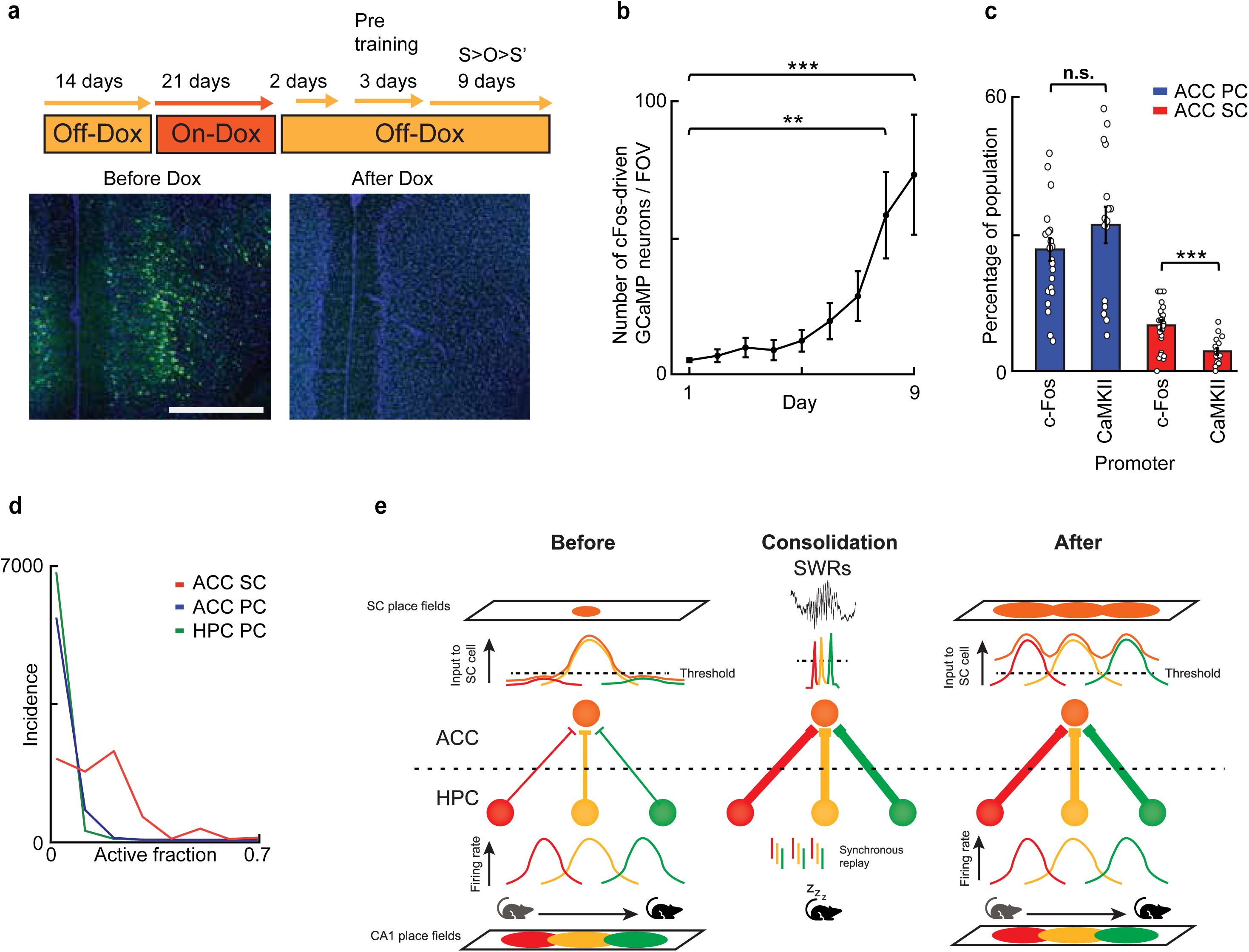
Spatial context cells form a synchronized cortical engram in ACC. (**a**) Experimental protocol and representative expression of c-Fos promoter mediated G-CaMP7 labelling. Scale bar = 500µm (**b**) Number of active neurons across days 1–9 from 6 mice. Friedman test (χ²(8) = 41.57, p < 0.0001), followed by Dunn’s multiple comparisons test relative to day 1. Significant increases were observed on day 8 (p = 0.0033) and day 9 (p = 0.0003). (**c**) Proportion of ACC place cells (PC, blue) and spatial context cells (SC, red) in *c-Fos*–tTA (5 mice) and TRE-GCaMP7 x CaMKIIα–tTA mice (4 mice). The proportion of place cells between *c-Fos* and CaMKIIα groups was comparable (Mann–Whitney U = 142, p = 0.1716), but the proportion of spatial context cells was significantly higher in *c-Fos*–tTA mice (Mann–Whitney U = 71.50, p = 0.0006). (**d**) Distribution of the co-active fraction of each neuron subclass during context O exploration on day 9. (**e**) A circuit-level model illustrating how hippocampal-cortical plasticity leads to the emergence of broad ACC receptive fields from discrete hippocampal (HPC) inputs. In a novel environment, an ACC neuron receives input from a specific CA1 cell assembly with low initial synaptic weights. Consequently, the SC cell only reaches suprathreshold firing at the center of individual CA1 place fields where input is maximal, resulting in sparse, multi-field activity (left). During offline periods (middle), synchronous replay of the hippocampal assembly during SWRs induces long-term potentiation (LTP). This repetitive co-activation strengthens the constellation of hippocampal-cortical synapses. In a familiar environment (right), the increased synaptic efficacy lowers the effective firing threshold for the SC cell, allowing it to integrate inputs across the entire spatial span of the assembly. This process transforms discrete hippocampal place fields into a broadened, continuous spatial representation in the ACC.

### Spatial context cells require hippocampal activity for their formation but not once they are formed

Hippocampus is essential for the initial formation of episodic memories which occurs rapidly^13,15–19^. In contrast, the formation of spatial context cells in ACC occurs with a temporal delay, raising the question of whether these processes are linked, or occur independently. To address this, hippocampal neuronal activity was inhibited by administering C21 to mice expressing hM4Di bilaterally along the dorsoventral axis of the hippocampus, 20 minutes prior to each of the nine daily S→O→S’ sessions (**Fig. 3c**). The fraction of both ACC place and spatial context cells in the control groups (hM4Di-expressing mice receiving a saline injection and naïve mice treated with C21) increased during the 9-day training period. By contrast, the fraction in C21-treated hM4Di-expressing mice was significantly lower **(Fig. 3d, e**). Furthermore, across-day rate-map correlations were significantly lower in the C21 group, demonstrating that hippocampal activity is essential for establishing a stable spatial map in ACC (**Fig. 3f**).

Once memories are consolidated, cortical regions (including ACC) become the site for long-term storage, facilitating recall in some cases independently of hippocampus^20–22^. To test if hippocampal output is required for the recall of a contextual representation already consolidated to ACC, C21 was administered on day 10, following 9 days of training with an intact hippocampus (**Fig. 3c)**. Despite the acute suppression of hippocampal output on Day 10, the fractions of ACC place and spatial context cells recruited were comparable to previous days (**Fig. 3g, h**). Moreover, both field size and across-session correlation (between days 8 × 9 vs 9 × 10) of spatial context cells were unaffected when hippocampal activity was suppressed on day 10 (**Fig. 3i, j**). This aligns with the memory consolidation theory, suggesting that the initial formation of a stable ACC contextual representation requires hippocampal input, but its maintenance and recall do not.

### Preferential recruitment of spatial context cells into the ACC engram supports remote memory retrieval

The immediate early gene, *c-Fos*, has been used as a marker to define memory engram bearing neurons ^11,23,24^. Since the contextual component comprises an integral part of any memory, we hypothesized that spatial context cells may be enriched in the *c-Fos* labeled population of neurons. To label active ACC cells during the encoding of a context specific memory, *c-Fos*-tTA transgenic mice were injected with AAV9-TRE-G-CaMP7 in ACC (**Fig. 4a**). After stopping doxycycline, we observed an increase in the number of G-CaMP7 labelled cells (**Fig. 4b**) and by days 8 and 9, 9.9±1.1% of all G-CaMP7 labelled neurons were spatial context cells, significantly higher than with CaMKIIα-promoter driven expressions (Mann–Whitney U = 71.50, p = 0.0006) (**Fig. 4c**). In contrast, the proportion of place cells was not significantly different between the two labeling strategies (**Fig. 4c**). These results indicate that the ACC spatial context cell may represent cortical memory engram.

Finally, to compare how focally the maps residing in hippocampus and ACC are activated during exploration, the fraction of co-active neurons was calculated across all periods of movement in context O (in 150 ms bins on day 9). The spatial coactivation dynamics (**Fig. 4e**) were similar between place cells in the hippocampus and ACC (0.06 vs 0.07; Ranksum N=5,4, p>0.05), suggesting equivalent levels of ensemble engagement. However, spatial context cells exhibited a significantly greater degree of co-activity (0.21; Ranksum N=5,4 p<0.05). To control for the inherent bias toward increased co-activity in cells with large receptive fields, we computed an expected co-activity score using a spatial shuffling procedure, which demonstrated that the observed synchrony among spatial context cells was significantly greater than predicted by their individual firing rates and receptive field size alone (**Extended Data Fig. 3**). This suggests the presence of a local circuit mechanism that selectively synchronizes the spatial context cells. Such synchronicity can effectively activate further downstream brain regions, or serve as self-perpetuating mechanism.

## Discussion

Here we attempted to identify the downstream effect of SWRs that occur during offline periods by employing closed-loop disruption and *in vivo* Ca^2+^ imaging. As a result, we identified a previously undescribed class of neuron in the anterior cingulate cortex (ACC), which we term spatial context cells, as the specific class of neurons generated by SWRs. These neurons are defined by activity patterns that span extensive portions of a specific environment while remaining largely silent or remapping in others (**Fig. 1–2**). Unlike place cells in the hippocampus or ACC, spatial context cells exhibit markedly broader receptive fields, with recruitment increasing progressively with repeated environmental exposure. This experience-dependent emergence reflects a transformation of their activity: they transition from initially fragmented, multi-field, place-like firing to a stable, single field encompassing a majority of the familiar environment **(Fig. 2**). Our results strongly support the idea that this neuronal subpopulation represents a cortical engram encoding specific contexts. Consistent with an engram function, these neurons are selectively enriched in the *c-Fos* labeled fraction after learning a novel task (**Fig. 4**). Moreover, their emergence and refinement require experience and adhere to the established timeline for memory consolidation from the hippocampus to cortical structures (**Fig. 3**).

We observed no anatomical segregation between ACC place and spatial context cells, suggesting they coexist as a heterogeneous population. While our data suggest that spatial context cells are likely predetermined by intrinsic connectivity rather than random assignment, the SWR disruption experiments (**Fig. 1 and 4**) coupled with the pattern of spatial context cell development (**Fig. 3**) we observed, point to a mechanism whereby highly convergent inputs from multiple dCA1 place cells, collectively spanning a large area, are consolidated as contextual memory traces in ACC neurons during SWR events (**Fig. 4f**). Given that spatial context cells follow a distinct multi-day developmental trajectory where initially fragmented, multi-field activity converges into singular, global representations, our data suggest that prior to consolidation, hippocampal inputs to the ACC are largely subthreshold, reaching the firing threshold only at their peak intensities. This results in a sparse ACC firing pattern characterized by multiple, restricted receptive fields. During offline states, however, SWR-mediated replay induces LTP in ACC neurons where these inputs converge, boosting previously subthreshold signals to suprathreshold levels. This synaptic potentiation allows the ACC firing field to expand, integrating locations that were initially represented only by weak hippocampal inputs. While our model focuses on the hippocampal-ACC axis, we cannot exclude the possibility that this convergence occurs within an intermediary region bridging the two structures.

The coexistence and differential engagement of spatial codes in the hippocampus and ACC are of particular interest. The hippocampal map is locally activated, with small receptive fields tracking the animal’s position. While SWRs and theta sequences briefly code non-local representations ^25,26^, these remain limited to spatially constrained areas or well-defined trajectories. We found that the ACC contains an almost identical population of place cells, with similar field sizes, stability, and local activation during exploration, though a high proportion of these neurons also conjunctively code for head-direction and likely other sensory information. In contrast to these localized spatial maps, ACC spatial context cells exhibit a qualitatively different engagement pattern. Since their receptive field encompasses, on average, over half of the environment, coupled with a high degree of co-activation (>20% of the population active simultaneously), these neurons comprise a globally activated code for context, which contrasts sharply with the local hippocampal code from which they are likely derived (**Fig. 4**). This high degree of co-activity and broad spatial coverage is not a computational artifact but rather a necessary functional property of the global contextual code generated by spatial context cells. Since the retrieval of a consolidated memory trace requires the near-simultaneous activation of an ensemble representing the entire environment, the high synchrony and expansive receptive fields of spatial context cells are optimized to provide this stable, non-localized contextual signal.

Our findings establish that the formation of the ACC spatial representation is dependent on hippocampal activity, as silencing the hippocampus disrupts the emergence and stability of both ACC place and spatial context cells **(Fig. 3-4**). What remains less clear is the precise anatomical route of this information transfer. While direct connectivity from the ventral hippocampus (vCA1) to the ACC is anatomically supported ^27–31^, evidence for direct projections from dorsal hippocampus (dCA1) is weak ^32–35^. This suggests an intermediate structure, such as the retrosplenial cortex (RSC) ^36,37^, may be necessary to convey dCA1 spatial information to the ACC. Given that the RSC contains spatially tuned, place-like neurons ^36,38,39^ and is ideally positioned to bridge these two structures, future studies targeting the RSC should clarify its role in sculpting the ACC spatial code.

In summary, this study identifies a novel class of ACC neuron that houses a long-term memory trace of context as the consequence of SWR/replay events. As a population, these neurons develop and mature with experience in a time-frame consistent with memory consolidation. Their expression relies on information routed from the hippocampus, specifically during SWRs, which appear to transform precise spatial information into a more generalized, stable, and abstractive cortical contextual representation. While their role in remote memory recall is clear, it remains an open question whether these context-bearing neurons themselves represent a complete memory engram or if they form part of a broader network encoding multiple features of specific episodes. Future work will undoubtedly address this question and further explore their therapeutic potential for disorders characterized by maladaptive contextual associations.

## Extended Data Figures

**Extended Data Fig. 1:**
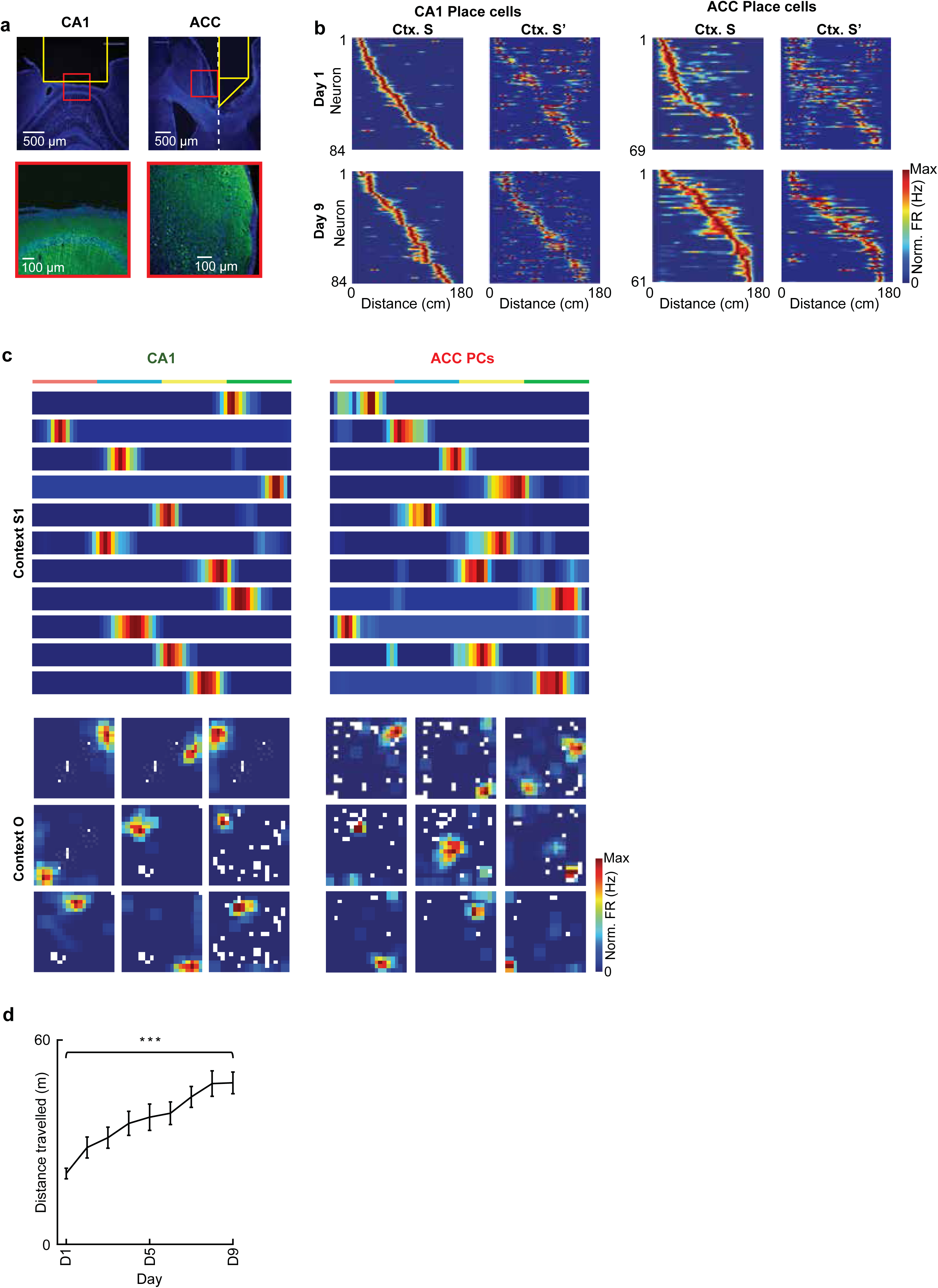
Place activity in dCA1 and ACC. (**a**) Representative coronal sections showing GCaMP7 fluorescence (green) and nuclear counterstaining with DAPI (blue) in dorsal CA1 (left) and anterior cingulate cortex (right). Yellow outlines indicate the approximate imaging area beneath the implanted GRIN lens, and red boxes denote the magnified regions shown below. Scale bars, 500 µm (top) and 100 µm (bottom). (**b**) Representative rate maps of CA1 and ACC place cells across sessions in a day. Each row shows the normalized firing rate of a single neuron, sorted by peak position in context. S. (**c**) Representative activity maps of dCA1 and ACC neurons during exploration of context S (top) and an open field (bottom). Examples show normalized firing rate maps of CA1 place cells and ACC place cells (n = 11 neurons, square track; n = 9 neurons, open field). Color scale represents normalized firing rate (from 0 to maximum). White areas indicate regions the mouse did not visit during the session. (**d**) Average distance travelled within the square track across 9 consecutive training days (n = 9 mice). One-way ANOVA, F(8, 153)= 7.09, Dunnett’s post hoc test, Day 1 vs Day 9, p < 0.0001.

**Extended Data Fig. 2:**
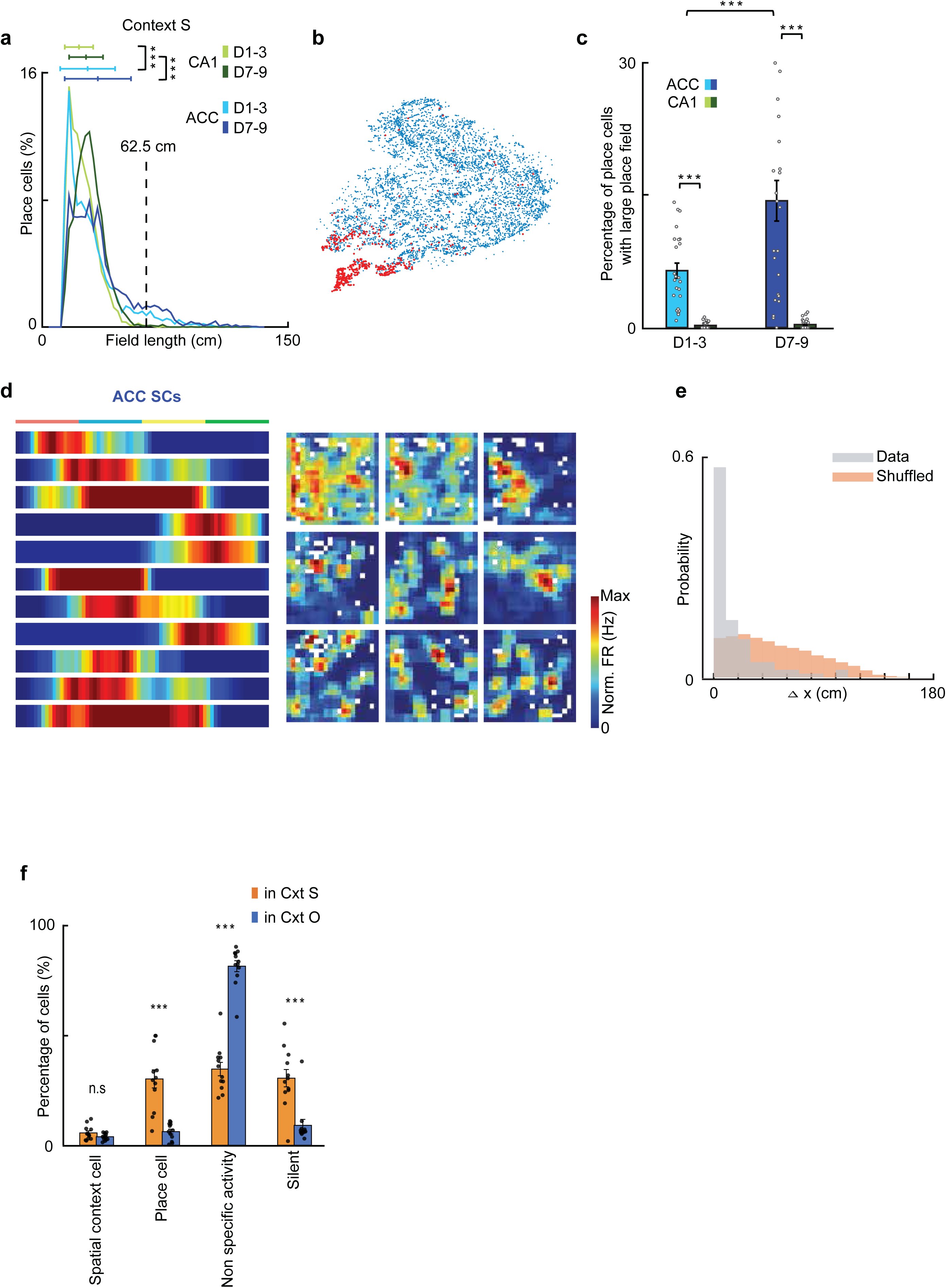
Identification of spatial context cells. (**a**) Distribution of place field size during early (D1–3) and late (D7–9) sessions in context S. ACC: n = 2452 neurons (4 mice, early), 2971 neurons (4 mice, late). CA1: n = 5161 neurons (5 mice, early), 3842 neurons (5 mice, late). Two-way ANOVA (Region × Phase): Region, F(1, 14425) = 693.77, P = 2.25 × 10⁻¹⁴⁹; Phase, F(1, 14425) = 480.45, P = 8.70 × 10⁻¹⁰⁵; Region × Phase, F(1,14425) = 16.57, P = 4.72 × 10⁻⁵. Post hoc Tukey–Kramer test: Early vs. Late, ACC (P < 0.0001), CA1 (P < 0.0001); ACC vs. CA1, early (P < 0.0001), late (P < 0.0001). Horizontal bars indicate mean ± SD. (**b**) Distribution of ACC neurons with place field widths exceeding 62.5cm (shown in red) within the t-SNE manifold. (**c**) The percentage of place cells with large place fields (≥62.5 cm) in the ACC (blue) and CA1 (green) during early (D1–3) and late (D7–9) phases. Each circle represents data from a single session (ACC: n = 24 sessions from 4 mice; CA1: n = 30 sessions from 5 mice). Two-way mixed-effects ANOVA with factors region (ACC vs. CA1) and period (early vs. late) revealed significant main effects of region (F(1, 11.27) = 21.78, P = 0.00064) and period (F(1, 99) = 25.05, P = 2.48 × 10⁻⁶), as well as a significant region × period interaction (F(1, 99) = 12.68, P = 0.00057). Post hoc Holm-adjusted pairwise comparisons showed that the fraction of large place fields was significantly greater in ACC than in CA1 during both early and late phases (early: P = 1.83 × 10⁻⁵, t = 4.67; late: P = 1.05 × 10⁻⁹, t = 6.99). Within-region comparisons indicated that the proportion increased from early to late sessions in ACC (P = 6.85 × 10⁻⁶, t = 5.00) but not in CA1 (P = 0.80, t = 0.25). (**d**) Additional examples of spatial context cell activity in the square linear track (left) and open field (right). (**e**) Histogram showing the shift in field centers (Δ centroid position) between neurons that were classified as spatial context fields in session S and subsequently place cells in session S′. Gray bars show the actual Δ centroid position distribution, while shuffled controls (orange) correspond to randomly circular shifted S′ rate maps. Δ centroid position values close to 0 indicate spatial stability, whereas higher values reflect remapping of the receptive field position. A significant difference was observed between the real and shuffled datasets (p < 1 × 10⁻²⁷, Wilcoxon rank-sum test). (**f**) Proportions of spatial context cells (SC), place cells (PC), non-specific activity cells (Other), and silent cells were quantified in familiar spatial contexts S and O using calcium imaging data collected on days 7–9 (4 mice × 3 days; n = 12 session-level datasets). Two-way ANOVA with Šidák’s multiple comparisons; F(3, 88) = 86.77, P < 0.0001), main effect of cell category (F(3, 88) = 176.2, P < 0.0001), main effect of context (F(1, 88) = 1.042e−014, P > 0.9999. Spatial context cell: p= 0.9981.

**Extended Data Fig. 3:**
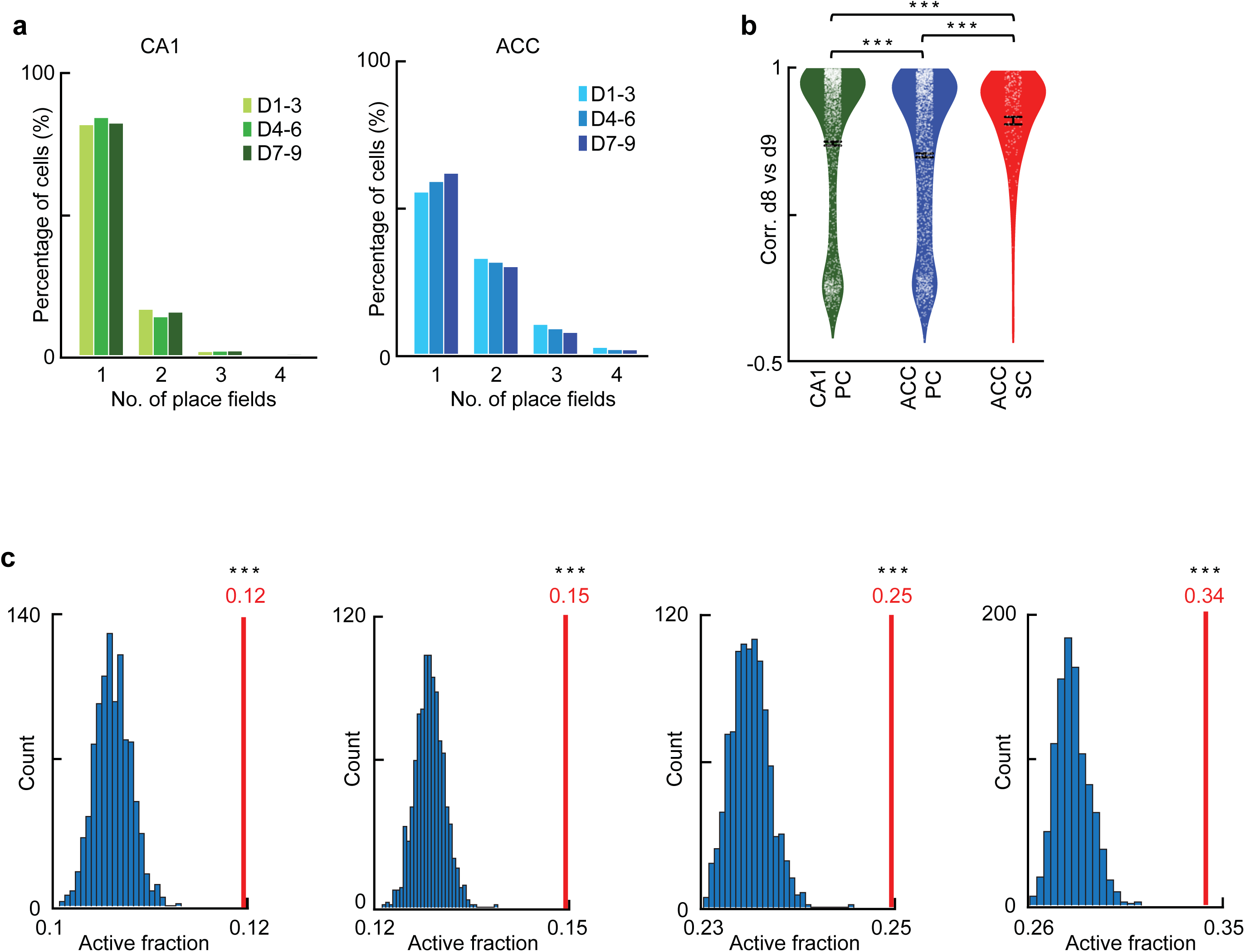
Additional properties of spatial context cells. (**a**) Histograms showing the proportion of neurons exhibiting 1–4 distinct place fields during the linear track task. Left, CA1; right, ACC. CA1 neurons predominantly had a single place field throughout the task period (D1–3, D4–6, D7–9), whereas ACC neurons often exhibited multiple spatial fields, particularly during later phases (D7–9). (**b**) Stability of spatial representations (correlation between D8 and D9 activity maps) for CA1 place cells (n = 1607 neurons, 10 sessions from 5 mice), ACC place cells (n = 1564 neurons, 8 sessions from 4 mice), and ACC SC cells (n = 212 neurons, 8 sessions from 4 mice). One-way ANOVA revealed a significant main effect of group (F(2, 3380) = 21.51, P < 0.0001). Tukey’s multiple comparisons test showed that ACC-SC cells exhibited significantly higher correlation values than both ACC-PC (mean diff = 0.1763, adjusted P < 0.0001) and CA1-PC (mean diff = 0.1158, adjusted P = 0.0003). In addition, CA1-PC cells showed significantly higher correlation values than ACC-PC (mean diff = 0.0606, adjusted P < 0.0001) (**c**) The observed active fractions of spatial context cells from each mouse (vertical red lines), compared with randomly shuffled distributions (1000 iterations) indicating their increased synchrony is significantly higher than expected for their field sizes and firing rates (Two-tailed permutation tests,P < 0.001)

## Methods

### Animals

We used TRE-G-CaMP7-T2A-DsRed2 x CaMKIIα-tTA double-transgenic mice to co-express GCaMP7 and DsRed2 selectively in excitatory neurons ^40^. This allowed for chronic, longitudinal observation of activity. In a subset of experiments, c-Fos-tTA mice ^41^, were used for activity-dependent labelling and chemogenetic manipulation.

### Subjects and housing

All procedures were conducted in strict accordance with institutional guidelines and protocols approved by the RIKEN and Kyoto University Animal Experiment Committees. Experiments utilized male mice (10–24 weeks old), which were housed individually on a reversed 12-hour light-dark cycle (lights on at 6 PM, off at 6 AM). All behavioral and imaging experiments were performed during the animals’ subjective dark phase (between 6 AM and 6 PM).

### Surgical procedures

All surgical procedures were conducted under isoflurane anaesthesia (3% induction, 1.5% maintenance) with the animal secured in a stereotaxic frame (Kopf).

#### Surgery for dorsal CA1 imaging

Mice were fixed in the stereotaxic frame, and a custom stainless-steel headplate (25 mm length, 4 mm width, 1 mm thickness) was affixed to the skull with dental cement and three anchor screws. The headplate featured a circular opening (7 mm inner diameter, 10 mm outer diameter) offset 2.5 mm relative to the plate center. The center of this opening was positioned at 2 mm AP, 2 mm ML from Bregma, targeting the left hemisphere. Three to four days post-headplate implantation, a trephine drill was used to remove a circular section of skull centered at the target coordinates. The dura and overlying cortex above CA1 were slowly removed by aspiration using a 27-gauge blunt needle, with exposed tissue continuously rinsed with cortex buffer (123 mM NaCl, 5 mM KCl, 10 mM glucose, 2 mM, 2 mM & 10 mM HEPES with a pH of 7.4). Finally, a metal guide tube was inserted into the cavity, positioned 200-300 µm superficial to the CA1 cell body layer, and secured directly to the skull using dental acrylic.

#### Surgery for ACC imaging

A stainless-steel headplate (identical to the hippocampal surgery) and four anchor screws were affixed to the skull with dental cement. Three to four days later, a circular craniotomy (2 mm diameter) was performed over the ACC. The opening was centered on the midline at 1 mm AP from Bregma, targeting the ACC. The dura overlying the implantation hemisphere was removed to access the longitudinal fissure. The microprism assembly, comprising a right-angle microprism bonded to a GRIN lens (0.85 mm diameter, 3.3 mm length or 9 mm length, Inscopix), was aligned parallel to the longitudinal fissure and positioned over the midline at 1 mm AP. The prism tip was used to gently retract the superior sagittal sinus (∼150 μm laterally) to facilitate unimpeded insertion. The microprism was slowly lowered into the subdural space within the longitudinal fissure to a final depth of 1.0 mm (from the cortical surface to the prism’s tip,), such that its front face was flush against the medial wall of the contralateral hemisphere, facing the left ACC. The craniotomy and any exposed brain tissue were sealed with silicone adhesive (Kwik-Sil), followed by a final application of dental cement to secure the microprism assembly to the skull ^42^.

For both CA1 and ACC imaging, the integrated miniature microscope (nVista system) was carefully lowered toward the respective lens/microprism assembly until the GCaMP7 fluorescence signal became clearly visible under the microscope’s LED light source (0.12–0.24 mW). After identifying a suitable field-of-view, the microscope’s base plate was fixed securely to the cranium using dental acrylic mixed with black acrylic paint to minimize light leakage. Before each daily behavioural experiment, the microscope was re-affixed to its base plate under brief isoflurane anaesthesia. Mice were allowed to fully recover in their home cage for 15 minutes prior to behavioural testing.

### Behavioural task and protocol

Mice performed a spatial navigation task across nine consecutive days, with each daily session consisting of three 10-minute epochs separated by 5-minute intermissions (**Fig. 1c**). These consisted of context S (Square Track): A square track (50 cm per edge) with a wall blocking one corner. Mice ran back-and-forth between the two open ends to receive food rewards, provided upon reaching each end. Performance improved over days, characterized by increased travelling distance, indicating familiarity **(Extended Data Fig. 1**). Context O (Open Field): A 50 x 50 cm open arena featuring distinct wall patterns and a unique scent (0.5% acetic acid). Food rewards were randomly scattered within the arena to promote general navigation. Finally, context S’: Mice were returned to the original square track (Context S). Mice were pre-exposed to the behavioral apparatus for 3 days which constituted the pretraining period. This consisted of one day in which mice could freely explore contexts S and O for 5 minutes. Followed by two days in which sucrose pellets (#1811251 Sucrose Rewards Tablet, Test Diet) were delivered at the ends of the square track and scattered randomly throughout context O. Throughout both pre-training and the experimental protocol, mice were moderately food deprived to maintain motivation to explore the contexts sufficiently. Immediately prior to the beginning of each session, tracks were wiped with scented paper towels (0.5% acetic acid for context O and 80% ethanol for contexts S & S’). The S→O→S’ protocol was repeated for a total of 9 consecutive days.

### High contextual overlap experiments

Mice were sequentially trained in three distinct square environments (contexts S_A_, S_B_ and S_C_), which had the same geometry and size as those used in the behavioral task but differed solely in sensory cues, including floor texture, wall texture, odour and background sound. Context S_A_ consisted of a smooth floor, fluffy wall texture, ethanol (EtOH) odour and continuous white noise. Context S_B_ contained a groovy floor, smooth plastic walls, acetic acid odour, and baroque background music. Context S_C_ was composed of a white Kim towel floor, metallic walls, almond odour and intermittent fan noise. Through days 1–8, animals were repeatedly exposed to contexts S_A_ followed by S_B_ (10 min per session), with a 5-min inter-session interval in their home cage. On day 9, mice followed the same protocol but were additionally exposed to the novel S_C_ context for 10 minutes. Calcium imaging was performed throughout all sessions under identical recording conditions.

### Viral injections

Surgical procedures were performed under isoflurane anesthesia (5% for induction, 1.5% for maintenance). Mice were placed in a stereotaxic apparatus (Kopf Instruments) and virus was injected using a glass micropipette attached to a microsyringe (MS-10; ITO) through a tube filled with liquid paraffin. A total of 150–500 nL of virus was injected at a rate of 60 nL/min. For experiments targeting broad hippocampal suppression, virus was injected at three sites per hemisphere to cover the rostrocaudal extent of the hippocampus. The coordinates (from bregma) were as follows: (AP +2.0 mm, ML ±1.4 mm, DV −1.8 mm), (AP +2.5 mm, ML ±2.1 mm, DV −1.6 mm), and (AP +3.0 mm, ML ±3.25 mm, DV −4.0 mm). For anterior cingulate cortex (ACC), injections were targeted at the following coordinates: +0.8 mm AP, –0.3 mm ML, and –1.7 mm DV. Following each injection, the pipette was left in place for an additional 10 min prior to withdrawal to prevent backflow. Following surgery, the scalp was sutured and mice were allowed to recover on a heating pad before being returned to their home cages. At least 3 weeks were allowed for viral expression prior to imaging or behavioral experiments. Analgesics were administered postoperatively as needed. All injection sites were verified histologically at the conclusion of the experiments.

### Histology & immunohistochemistry

Following the conclusion of imaging experiments, mice were anesthetized and perfused transcardially with phosphate-buffered saline (PBS) followed by paraformaldehyde (PFA). Brains were extracted, post-fixed in PFA for 24 hours, and then transferred to PBS for an additional 24 hours. Coronal sections (50 µm thickness) were cut using a Leica vibratome. Sliced tissue sections were incubated overnight at in a buffer solution (0.1 M Tris-HCl, 0.15 M NaCl) containing 0.5% Triton-X and a 5% blocking reagent (Roche), along with the primary antibody (Rabbit anti-GFP antibody, Thermo Fisher Scientific, A11122, 1:500). Sections were then rinsed three times in PBS (15 minutes each) and incubated with the AlexaFluor 488-conjugated secondary antibody (Cell Signalling, 1:500). After a final three washes in PBS (15 minutes each), slices were mounted and coverslipped using mounting medium containing DAPI (Vector Laboratories). Images were captured using both confocal and standard fluorescent microscopes to confirm lens placement and protein expression.

### DREADD mediated inhibition

For chemogenetic silencing experiments, the DREADD agonist 21 (Cayman Chemical Company, #18907), an alternative to clozapine N-oxide (CNO) was used to activate the hM4D receptor. First, a 5mg/ml stock solution dissolved in DMSO was prepared, then diluted in saline to the desired concentration (0.05mg or 0.1mg/ml) prior to experiments. C21 was injected intraperitonially (i.p.) at a dose of 1mg/kg, 20 minutes prior to the start of behavioural experiments. For offline chemogenetic inhibition, mice received an intraperitoneal injection of C21 at 2 hours and 5 hours after each daily behavioral session during days 1–4.

### Doxycycline preparation

Doxycycline (Dox) powder (5 g; Thermo Fisher Scientific, Cat. No. 458890050) was dissolved in Milli-Q water to prepare a 50 mg/mL stock solution. This solution was aliquoted into in 15 mL centrifuge tubes (4 ml per tube) and stored at −20°C until use. Prior to injection, the stock solution was diluted with Milli-Q water to a final concentration of 2 mg/mL and provided ad libitum as drinking water. Water bottles were wrapped with aluminum foil, as doxycycline is known to be light sensitive.

### *c-Fos*–labeled neuron imaging

To enable activity-dependent calcium imaging, c-fos-tTA mice were injected with AAV9-TRE-G-CaMP7 into the ACC. Following viral injection, mice were maintained in the doxycycline (Dox)-OFF condition for 3 weeks to allow recovery and sufficient G-CaMP7 expression. Expression levels were confirmed by microscope-based imaging prior to behavioral experiments. After confirming that fluorescence signals were detectable and suitable for imaging, mice were administered Dox in their drinking water for 21 days to suppress expression. Dox was then withdrawn for 2 days, after which mice underwent 3 days of pre-training followed by the 9-day behavioral task described above.

### Calcium imaging data processing

Calcium imaging data were processed using Mosaic (Inscopix). To increase computation speed, images were spatially down-sampled by a factor of four in each dimension. Next, the imaging data were then motion corrected and converted to relative fluorescence changes (ΔF/F₀), defined as:

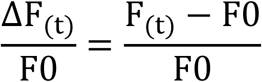

where F₀ represents the baseline fluorescence for each pixel, calculated as the mean fluorescence across the entire recording period. Finally, the movie was smoothed using a disk averaging filter (radius: 3 × 3 pixels). The filtered imaging data were used for neuron identification and clustering score calculation only, and not for fluorescence change analysis (see ‘neuron identification’ and ‘calcium event identification’).

### Neuron identification

Regions of interest (ROIs) corresponding to individual neurons were identified using a custom MATLAB script. For each motion-corrected frame, the image was converted to double precision and globally standardized by subtracting the frame-wise mean intensity. A binary activity map was then generated by thresholding pixels above the frame-wise mean plus SD × σ, where σ denotes the frame-wise standard deviation and SD is a user-defined sensitivity parameter. The binary image was multiplied by a predefined field-of-view mask, hole-filled, and segmented into 8-connected components. For each component, area and x/y spans were calculated from the pixel coordinates, and candidates were retained if their area was between 30 and 180 pixels (approximately 250–1500 µm²) and their width-to-height and height-to-width ratios were both < 2. Surviving masks from all frames were pooled and deduplicated based on spatial similarity. Specifically, cosine similarity between vectorized binary masks was computed, and when the similarity exceeded 0.4, the smaller mask was discarded and the larger mask was retained as the representative ROI. The resulting nonredundant set of ROI masks was used as the neuron list for each session.

### Calcium event identification

ΔF/F traces were extracted for each ROI from the un-filtered calcium imaging movie. To remove crosstalk between neighbouring ROIs, a clustering score for each ROI was calculated for every frame as follows. First, each ROI was enlarged 5 times maintaining the original center position. Second, the intensity of each pixel within this enlarged ROI was calculated and only the brightest 20% of pixels were kept. Third, the clustering score was calculated as the proportion of the brightest subset of pixels that were located in the original ROI. In this case, low clustering scores are obtained when activity in the original ROI is minimal, or when neighbouring neuron activity is higher than in the ROI of interest. Conversely, high clustering scores are obtained only when the cell in the original ROI fires. By excluding values with low clustering score from each ROI, crosstalk was minimized. These steps were repeated for each frame and each ROI. Calcium events were detected when both the smoothed clustering score exceeded 0.4 and the normalized fluorescence signal exceeded 1.54 standard deviations above baseline.

### Neuron tracking

Following alignment, ROIs across sessions were matched based on both spatial proximity and footprint similarity. For each ROI, candidate matches in other sessions were identified if their centroid distance was less than 4 pixels (∼10 µm) and the cosine similarity of their spatial footprints exceeded 0.6. When multiple candidate matches were found, the ROI with the highest similarity-to-distance ratio (similarity/distance) was selected as the corresponding neuron.

### Place cell classification

#### Open field

For place cell identification, spatial information was computed using a coarse spatial binning (3 × 3 bins, 16.7 × 16.7 cm / bin). Calcium events were assigned to bins based on the animal’s position, and only events during running periods (speed > 1 cm/s) were included. Event rates were calculated by dividing event counts by occupancy. Spatial information was computed as previously described (Markus et al., 1994)

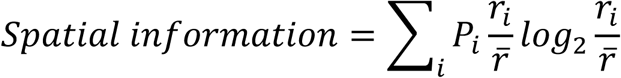

Where *r_i_* is the calcium event rate of the neuron in the *i^th^* bin, *p_i_* is the probability of the mouse being in the *i^th^* bin and *r* is the overall mean calcium event rate. Next, a shuffled distribution of spatial information values was calculated for each neuron, by randomly permuting calcium activity across time 1000 times, such that they were no longer aligned with animal position. By comparing the observed value to the shuffled distribution, a significance value was calculated for each neuron’s spatial information. Only neurons with significant spatial information (p < 0.05) and mean firing rates higher than 0.1 Hz were classified as place cells. To detect place fields, firing rate maps were generated with a 2.5 cm x 2.5 cm spatial bin size, then normalized by the maximum firing rate in the map and smoothed with a 1 SD Gaussian kernel. Next, maps were binarized such that bins with firing rates >0.2 were converted to 1, while other bins were assigned 0. Contiguous bins in this binary map were detected by using the bwlabel function (pixel connectivity= 8) in MATLAB. Each collection of connected bins was considered an individual place field, with the main place field being designated as that with the highest firing rate.

#### Square track

To identify place cells in the square track, we used a reliability score to quantify the consistency of a neuron to repeatedly fire in its preferred location on the track. First, the track was divided into 72 equally spaced bins (2.5cm each), with the first and last 5 bins, where food rewards were delivered and consumed excluded. Next, for each lap a binarized activity map was generated where 1s and 0s represented the presence and absence of calcium events, respectively. The similarity between lap-wise activity maps was quantified using the spike time tiling coefficient (STTC) between all pairs of laps within a session. The average of these values was defined as the reliability score. To assess significance, for each neuron the binarized activity maps were shuffled 1000 times to generate a shuffled distribution, with only neurons showing significant (p < 0.05) reliability being considered place cells in the square track. To detect place fields, firing rate maps were normalized by their maximum value, smoothed with a 1 SD Gaussian kernel, and binarized (spatial bin with firing rate > 0.2 = 1, other spatial bins = 0). Contiguous bins in this binary map were detected and classified as a place field, with the maximum firing rate containing field designated as the main place field. These criteria were applied independently to forward and backward directions

### Identification of spatial context cells

#### Open field

In the open-field environment, calcium event rate maps were computed across a 20 × 20 spatial bin matrix using running frames only (speed > 1 cm/s). Event-rate maps were Gaussian-smoothed (σ = 1) and normalized to their peak. The spatial coverage of each neuron was defined as the proportion of bins with non-zero activity relative to all occupied bins. Neurons with a spatial coverage greater than 25% were classified as spatial context cells. This criterion was applied independently of place-field formation and used to identify broadly distributed context-related activity.

#### Square track

To identify spatial context cells in the square track, spatial firing rate maps were peak-normalized after Gaussian smoothing. The main field size was defined as the number of contiguous bins around the peak with a normalized rate ≥ 0.2. The total number of active bins (smoothed rate > 0) and bins above a secondary threshold (≥ 0.1) were also quantified for descriptive purposes. Neurons were classified as spatial context cells if they exhibited a main field > 62.5 cm (25 bins) and a peak-normalized rate > 0.3, parameters that largely captured this population of neurons. These criteria were applied independently for forward and backward running directions.

### Population vector correlation

To determine the level of similarity between representations of different sessions, mean population vector correlations were calculated ^43^. For each spatial bin (excluding 5 bins at each end of the track) rate vectors were constructed for each neuron, as the mean event rate per spatial bin. The correlation between the population vectors of two sessions were computed for all neurons and averaged across spatial bins.

### Location decoding

To decode animal location from calcium transient activity, the real position data were first linearized and smoothed before being divided into 50 equally spaced bins (4 cm). Next, a random 50 percent subset of the calcium transient data and the corresponding animal location data were used to train a multiclass error-correcting output codes (ECOC) model using support vector machine (SVM) binary learners in Matlab (using the fitcecoc function). The remaining calcium data were then used to decode animal location using the trained model and the percentage error between the real and decoded positions were calculated as a fraction of the entire track’s length.

### Context decoding

To decode session identity between the square linear track and open field, the calcium data for both sessions (in a single day) were combined and labelled according to session. Next, a random 20 percent subset of the calcium transient data and the corresponding identity labels were used to train a binary linear classifier model in Matlab (using the fitclinear function). The remaining calcium data were then used to decode the session identity using the trained model, with the error being expressed as a percentage of the entire decoded dataset.

To decode session identity between the three similar square linear tracks which differed only in sensory cues, a similar approach was used whereby the three sessions were concatenated and labelled accordingly. A random 20 percent subset of the calcium transient data was then used to train a multiclass error-correcting output codes (ECOC) model using support vector machine (SVM) binary learners in Matlab (using the fitcecoc function). The remaining calcium data were then used to decode the session identity using the trained model, with the error being expressed as a percentage of the entire decoded dataset.

### Dimensionality reduction and manifold analysis

To characterize the spatial firing properties of ACC and dCA1 neurons, we first generated occupancy-normalized ratemaps for each neuron. Mouse position was smoothed using a 15-frame moving average, and instantaneous direction was calculated to categorize activity into forward and backward trajectories. For each direction, the track was discretized into spatial bins, and the firing rate per bin was calculated as the total number of calcium transients divided by the time spent in that bin. To ensure robust spatial tuning, we calculated a stability index by correlating rate maps from the first and second halves of each session; neurons were classified as directionally selective if their stability in one direction significantly exceeded the other, or if their peak firing rate in one direction was at least double that of the opposite direction. To visualize high-dimensional neural population activity across nine days of navigation, we applied t-distributed stochastic neighbor embedding (t-SNE). For initial spatial mapping, t-SNE was performed on the population of all recorded neurons, with the resulting two-dimensional coordinates color-coded by the center of each neuron’s place field. To isolate functional clusters independently of spatial topology, we performed a manifold analysis on auto-correlated ratemaps of the ACC population. By using the auto-correlation, we removed absolute spatial position while preserving the characteristic “width” and periodicity of each neuron’s firing pattern. The resulting manifold was then analyzed for local neighborhood enrichment to identify subpopulations specifically affected by SWR disruption. Specifically, for each neuron, we calculated the proportion of its 150 nearest neighbors that belonged to the SWR-disruption group. Clusters within the t-SNE manifold were identified based on local density and neighborhood composition. To determine if specific clusters were depleted of SWR-disrupted neurons, we performed a Chi-square test on a 3×2 contingency table (comparing control, stimulated, and delayed groups against their presence in the target cluster versus the remainder of the population). Further quantification of place field sizes was performed by thresholding the spatial ratemaps; a threshold of 62.5 cm was determined to optimally differentiate the “spatial context cell” subpopulation from canonical place cells.

### Sharp-wave ripple suppression

To suppress sharp-wave ripples recording and stimulating electrodes were constructed by twisting and heating coated stainless-steel wire (0.0045”, A-M systems) into bipolar electrodes. To record SWRs from the CA1 cell body layer, two bipolar electrodes were bonded with a small amount of cyanoacrylate glue and cut at an ∼60° angle so that the tips of the electrodes were staggered and spanned a depth of approximately 1mm. Following anaesthetization (of TRE-GaMP7 x CaMKⅡ-tTA double transgenic mice) with isoflurane as previously described, two small craniotomies were made at -1.82 mm AP, -1.5 mm ML and -0.82 mm AP, -0.75 mm ML, into which the recording electrode bundle and a single bipolar stimulating electrode were inserted, to depths of 1.25 mm and 1.75 mm respectively from the brain’s surface. A small amount of Vaseline was used to seal the exposed brain surface surrounding the electrodes, which were subsequently fixed in place with dental acrylic. Next, a craniotomy was made and a microprism (Proview Integrated Prism 9 mm long, 1mm diameter, Inscopix) was inserted as previously described to target ACC. Any surrounding brain tissue was covered with silicone adhesive (Kwik-Sil, World Precision Instruments), and the prism fixed in place with dental acrylic. Four additional skull screws were inserted ∼ 2.8mm posterior from bregma, where space allowed. Two of these served as a ground and reference, while the other two were simply to anchor the implants to the skull. The electrode interface board (EIB, Neuralynx) glued to a custom 3D-printed mount was held in place by a micromanipulator at the back of the skull and after the ground and reference wires were connected with the screws, all components were bonded with dental acrylic mixed with black pigment. Finally, the bipolar stimulating electrode and the four recording electrodes were connected to the EIB and secured with gold-plated pins, after which a plastic cone was glued behind the implant to protect it from physical damage during grooming.

Implanted animals were subjected to the same experimental protocol as previously described, allowed to freely explore the S→O→S’ contexts for nine successive days. Immediately after the end of the S’ session, animals were transferred to a small sleep box where they were allowed to rest for a period of 1 hour, during which SWRs were suppressed in real time (or with a 200 ms delay which served as the control group) by ventral hippocampal commissure stimulation. Wide-band LFP signals were filtered in the sharp wave ripple band (120-300 Hz) allowing for SWRs to be detected online, using the closed loop signal processing platform (Hardware Processing Platform, Neuralynx). Online detection of ripples (threshold crossing events of the bandpass-filtered signal) automatically triggered a single-pulse (0.5 ms) commissural stimulation. The amplitude of the biphasic electrical pulses varied from 50 to 250 μA and was set in each session to the lowest amplitude that resulted in consistent disruption of hippocampal ripple events. Following the nine-day stimulation protocol, animals continued to undergo the same experimental schedule (S→O→S’) for a further nine days in the absence of SWR suppression.

### Head Direction Tuning Analysis

The directional tuning of recorded neurons was analyzed using custom Matlab scripts. The nose, head and animal body were labelled and tracked using deeplabcut across behavioural sessions. The animal’s head direction (HD) was determined by computing the angle between the head position and the body position (center of mass), yielding an angle in degrees (ranging from 0° to 360°). Animal position data were smoothed and periods of low velocity were identified and excluded: still periods, defined as times when the absolute change in smoothed *x* and *y* position fell below a threshold velocity of 4 cm/s. Additionally, neurons with an insufficient number of calcium transients (<5) throughout the 10-minute session were also excluded from further analyses. For each remaining neuron, the heading angle at the precise time of each calcium transient was extracted, creating a list of spike-associated angles.

### Quantification and Statistical Testing: Mean Resultant Vector Length (MVL)

The strength of directional tuning was quantified using the Mean Resultant Vector Length (MVL). Spike-associated angles were grouped into 18 equally spaced bins (20° per bin) to create a circular histogram of firing rate versus HD. The MVL, which ranges from 0 (no tuning) to 1 (perfect tuning), was calculated as follows:

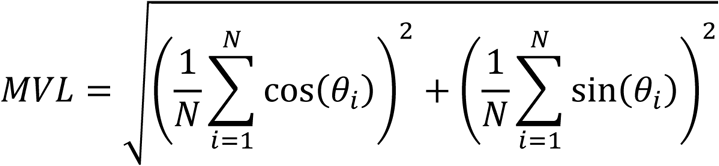

where *N* is the number of spikes and *θ*_*i*_ is the heading angle for the *i*-th spike.

### Shuffling Test for Significance

To determine the statistical significance of the MVL for each neuron, a shuffling test was performed. The observed MVL was compared to a distribution of MVLs generated from 500 shuffled iterations. In each iteration, a set of random heading angles was selected from the full session’s filtered HD, with the number of randomly sampled angles matching the neuron’s actual calcium transient count. The MVL was calculated for each shuffled dataset and the p-value defined as the proportion of shuffled MVLs that were greater than or equal to the observed MVL.

### Synchronization Analysis

Still periods were identified and excluded as described above, as were neurons with total spike counts <5. The remaining filtered transient data were stored in a matrix, where rows represent time steps (50 ms) and columns represent individual neurons. This matrix was padded with zeros at the end to ensure the total number of time steps (rows) was divisible by 3 and reshaped to group 3 consecutive time steps together. The resultant matrix was binarized (with 1 indicating a detected calcium transient), so that each row now corresponded to the activity of each neuron in 150 ms time steps. The fraction of simultaneously active neurons for each time bin was calculated by summing the number of active neurons (rows) in the final binned data and dividing by the total number of neurons. The overall synchronization value for the session was then computed as the mean of these non-zero co-activity fractions.

### Shuffling Test for Significance

To determine whether the increased synchrony of spatial context cells was statistically significant a shuffling test was performed. For each mouse, data were velocity thresholded as previously described. Next, the binarized firing maps for each spatial context cell were independently circularly permuted by a random amount, thus maintaining their firing rates and their spatial distribution across the open field. The fraction of simultaneously active neurons was then calculated and this process was repeated for 1000 iterations, with the p-value defined as the proportion of shuffled values that were greater than or equal to the observed fraction of simultaneously active neurons.

### Statistics

All statistical analyses were performed using GraphPad Prism 10 or MATLAB. Statistical tests, sample sizes, and definitions of significance are described in the corresponding figure legends. Data were tested using parametric or non-parametric tests as appropriate, including one-way, two-way, and mixed-effects ANOVA, as well as two-tailed paired or unpaired t-tests. Post hoc multiple comparisons were performed using appropriate corrections, as specified in the figure legends. Data are presented as mean ± SEM unless otherwise indicated.

## Resource availability

### Lead contact

Further information and requests for resources and reagents should be directed to, and will be fulfilled by, the lead contact, Yasunori Hayashi.

### Materials availability

Materials and reagents generated in this study will be made available upon request to the lead contact.

### Data and code availability

All code and data generated in this study will be made available on request.

## Acknowledgements

We thank Masamichi Ohkura and Junichi Nagai for G-CaMP7, Mark Mayford and Naoki Matsuo for providing the c-Fos-tTA mouse line, Thomas J. McHugh, Kenji Mizuseki and Shigeyoshi Fujisawa for their comments on the manuscript. This work was supported by RIKEN, Grant-in-Aid for Scientific Research, JP22110006, JP90466037, JP16H01292, JP19H01010, JP22H00434, JP22H04981, JP22K21353, JP22KF0211 from MEXT, The Uehara Memorial Foundation, The Naito Foundation, Research Foundation for Opto-Science and Technology, Novartis Foundation, and The Takeda Science Foundation, HFSP Research Grant RGP0022/2013, and JST CREST JPMJCR20E4 (Y.H.).

## Author contributions

Y.H. conceived the study. A.B., Y.K.M., S.J.M., X.J & S.T. performed the experiments. A.B., S.J.M., Y.K.M., S.T., Y.H., A.G. & A.L. analyzed the data. Y.H., T.T & S.J.M. supervised the project. A.B., S.J.M. & Y.H. wrote the manuscript.

## REFERENCES

1 Foster, D. J. & Wilson, M. A. Reverse replay of behavioural sequences in hippocampal place cells during the awake state. Nature 440, 680–683 (2006).

2 Diba, K. & Buzsaki, G. Forward and reverse hippocampal place-cell sequences during ripples. Nat. Neurosci. 10, 1241–1242 (2007).

3 Girardeau, G., Benchenane, K., Wiener, S. I., Buzsaki, G. & Zugaro, M. B. Selective suppression of hippocampal ripples impairs spatial memory. Nat. Neurosci. 12, 1222–1223 (2009).

4 Buzsaki, G. Hippocampal sharp wave-ripple: A cognitive biomarker for episodic memory and planning. Hippocampus 25, 1073–1188 (2015).

5 Kovacs, K. A. et al. Optogenetically Blocking Sharp Wave Ripple Events in Sleep Does Not Interfere with the Formation of Stable Spatial Representation in the CA1 Area of the Hippocampus. PLoS One 11, e0164675 (2016).

6 Norimoto, H. et al. Hippocampal ripples down-regulate synapses. Science 359, 1524–1527 (2018).

7 Singer, A. C., Carr, M. F., Karlsson, M. P. & Frank, L. M. Hippocampal SWR activity predicts correct decisions during the initial learning of an alternation task. Neuron 77, 1163–1173 (2013).

8 McClelland, J. L., McNaughton, B. L. & O’Reilly, R. C. Why there are complementary learning systems in the hippocampus and neocortex: insights from the successes and failures of connectionist models of learning and memory. Psychol Rev 102, 419–457 (1995).

9 Tse, D. et al. Schemas and memory consolidation. Science 316, 76–82 (2007).

10 Ego-Stengel, V. & Wilson, M. A. Disruption of ripple-associated hippocampal activity during rest impairs spatial learning in the rat. Hippocampus 20, 1–10 (2010).

11 Frankland, P. W., Bontempi, B., Talton, L. E., Kaczmarek, L. & Silva, A. J. The involvement of the anterior cingulate cortex in remote contextual fear memory. Science 304, 881–883 (2004).

12 O’Keefe, J. & Dostrovsky, J. The hippocampus as a spatial map. Preliminary evidence from unit activity in the freely-moving rat. Brain Res 34, 171–175 (1971).

13 Eichenbaum, H. Prefrontal-hippocampal interactions in episodic memory. Nat Rev Neurosci 18, 547–558 (2017).

14 Taube, J. S. Head direction cells and the neurophysiological basis for a sense of direction. Prog Neurobiol 55, 225–256 (1998).

15 Chandra, S., Sharma, S., Chaudhuri, R. & Fiete, I. Episodic and associative memory from spatial scaffolds in the hippocampus. Nature 638, 739–751 (2025).

16 Chettih, S. N., Mackevicius, E. L., Hale, S. & Aronov, D. Barcoding of episodic memories in the hippocampus of a food-caching bird. Cell 187, 1922–1935 e1920 (2024).

17 Dong, C., Madar, A. D. & Sheffield, M. E. J. Distinct place cell dynamics in CA1 and CA3 encode experience in new environments. Nat Commun 12, 2977 (2021).

18 Hill, A. J. First occurrence of hippocampal spatial firing in a new environment. Exp Neurol 62, 282–297 (1978).

19 Kolibius, L. D. et al. Hippocampal neurons code individual episodic memories in humans. Nat Hum Behav 7, 1968–1979 (2023).

20 Varela, C. et al. Tracking the Time-Dependent Role of the Hippocampus in Memory Recall Using DREADDs. PLoS One 11, e0154374 (2016).

21 Wiltgen, B. J. et al. The hippocampus plays a selective role in the retrieval of detailed contextual memories. Curr Biol 20, 1336–1344 (2010).

22 Bayley, P. J., Hopkins, R. O. & Squire, L. R. Successful recollection of remote autobiographical memories by amnesic patients with medial temporal lobe lesions. Neuron 38, 135–144 (2003).

23 Liu, X. et al. Optogenetic stimulation of a hippocampal engram activates fear memory recall. Nature 484, 381–385 (2012).

24 Kitamura, T. et al. Engrams and circuits crucial for systems consolidation of a memory. Science 356, 73–78 (2017).

25 Middleton, S. J. & McHugh, T. J. Silencing CA3 disrupts temporal coding in the CA1 ensemble. Nat Neurosci 19, 945–951 (2016).

26 Middleton, S. J. et al. Altered hippocampal replay is associated with memory impairment in mice heterozygous for the Scn2a gene. Nat Neurosci 21, 996–1003 (2018).

27 Degenetais, E., Thierry, A. M., Glowinski, J. & Gioanni, Y. Synaptic influence of hippocampus on pyramidal cells of the rat prefrontal cortex: an in vivo intracellular recording study. Cereb Cortex 13, 782–792 (2003).

28 Dembrow, N. C., Chitwood, R. A. & Johnston, D. Projection-specific neuromodulation of medial prefrontal cortex neurons. J Neurosci 30, 16922–16937 (2010).

29 Liu, X. & Carter, A. G. Ventral Hippocampal Inputs Preferentially Drive Corticocortical Neurons in the Infralimbic Prefrontal Cortex. J Neurosci 38, 7351–7363 (2018).

30 Phillips, M. L., Robinson, H. A. & Pozzo-Miller, L. Ventral hippocampal projections to the medial prefrontal cortex regulate social memory. Elife 8 (2019).

31 Thierry, A. M., Gioanni, Y., Degenetais, E. & Glowinski, J. Hippocampo-prefrontal cortex pathway: anatomical and electrophysiological characteristics. Hippocampus 10, 411–419 (2000).

32 Andrianova, L. et al. No evidence from complementary data sources of a direct glutamatergic projection from the mouse anterior cingulate area to the hippocampal formation. Elife 12 (2023).

33 Glat, M. et al. An accessory prefrontal cortex-thalamus circuit sculpts maternal behavior in virgin female mice. EMBO J 41, e111648 (2022).

34 Rajasethupathy, P. et al. Projections from neocortex mediate top-down control of memory retrieval. Nature 526, 653–659 (2015).

35 Shi, W. et al. Whole-brain mapping of efferent projections of the anterior cingulate cortex in adult male mice. Mol Pain 18, 17448069221094529 (2022).

36 Gianatti, M. et al. Multiple long-range projections convey position information to the agranular retrosplenial cortex. Cell Rep 42, 113109 (2023).

37 Sugar, J., Witter, M. P., van Strien, N. M. & Cappaert, N. L. The retrosplenial cortex: intrinsic connectivity and connections with the (para)hippocampal region in the rat. An interactive connectome. Front Neuroinform 5, 7 (2011).

38 Mao, D., Kandler, S., McNaughton, B. L. & Bonin, V. Sparse orthogonal population representation of spatial context in the retrosplenial cortex. Nat Commun 8, 243 (2017).

39 Sun, H. et al. Conjunctive processing of spatial border and locomotion in retrosplenial cortex during spatial navigation. J Physiol 602, 5017–5038 (2024).

40 Sato, M., et al. Generation and Imaging of Transgenic Mice that Express G-CaMP7 under a Tetracycline Response Element. PLoS One 10, e0125354 (2015).

41 Matsuo, N., Reijmers, L. & Mayford, M. Spine-type-specific recruitment of newly synthesized AMPA receptors with learning. Science 319, 1104–1107 (2008).

42 Low, R. J., Gu, Y. & Tank, D. W. Cellular resolution optical access to brain regions in fissures: imaging medial prefrontal cortex and grid cells in entorhinal cortex. Proc Natl Acad Sci U S A 111, 18739–18744 (2014).

43 Leutgeb, J. K. et al. Progressive transformation of hippocampal neuronal representations in “morphed” environments. Neuron 48, 345–358 (2005).

